# Attractive and repulsive history effects in categorical and continuous estimates of orientation perception

**DOI:** 10.1101/2025.04.24.650437

**Authors:** Mert Can, Thérèse Collins

## Abstract

Perceptual reports can be attracted towards or repulsed from previous stimuli and responses. We investigated the conditions in which attractive and repulsive history effects occur with oriented Gabors by manipulating response type and frequency, and stimulus duration. When subjects adjusted a continuous response cue to match orientation, we observed repulsion from the previous stimulus when the stimulus was presented for 50 ms and attraction from the previous stimulus and response when it was presented for 500 ms, regardless of whether responses were given to every stimulus or every other stimulus. With a categorical clockwise/counter-clockwise response, there was attraction to the previous response and repulsion from the previous stimulus. Attraction to the previous response was stronger with sequential responses and short relative orientations. Repulsion was constant across all stimulus durations and response frequencies, and increased with relative orientation. The overall history effect of the previous response and stimulus was repulsive with alternating categorical responses and attractive with sequential categorical responses. Our results replicate and synthesize seminal findings of the serial dependence and adaptation literature, and show independent history effects working with and against each other determined by whether the response is categorical or continuous.

## Introduction

The human brain leverages redundancies in the visual world to extract information from it despite the constant influx of input. Integrating information across time may reduce the amount of spurious variations selected for further processing, but this may come with a cost. One of these costs is an erroneous perception of objects or their features.

The integration of information across time can be studied via history effects, such as adaptation (when exposure to one visual stimulus alters the perception of the next stimulus) or serial dependence (when consecutively presented stimuli are seen to resemble each other). The erroneous perception caused by these history effects can be measured in reports about the visual features of objects. For example, a vertically oriented Gabor patch could be reported to be more counter-clockwise (CCW) or clockwise (CW) depending on a previous Gabor’s orientation. In other words, the current stimulus may be attracted towards or repulsed away from the previous stimulus. Attractive effects may reveal how our visual system acts on temporal redundancies between consecutive objects to provide visual stability across time, and repulsion may reveal how the visual system acts on informational redundancies in an object to increase sensitivity for novelty. Ideally, observers could switch flexibly between attraction and repulsion as a function of task demands.

In recent years, there has been increased interest in understanding history effects, in particular the attractive effect called serial dependence (Fischer & Whitney, 2014). Attentional, temporal, spatial, and featural factors are known to facilitate serial dependence (for review, see Manassi, Munai, & Whitney, 2023; Pascucci, Tanrikulu, Ozkirli, Houborg, Ceylan, Zerr, Rafiei, & Kristjánsson, 2023), however, there is little agreement about the nature of serial dependence (SD). SD could be a response bias or decisional effect, subjects repeating their previous responses when it is difficult to rely on the current stimulus percept (Bosch, Fritsche, Utzerath, Buitelaar, & de Lange, 2022; Schwiedrzik, Ruff, Lazar, Leitner, Singer, & Melloni, 2014; Van Geert, Moors, Haaf, & Wagemans, 2022), or stimuli facilitating biases in decisional stages of visual processing (Pascucci, Mancuso, Santandrea, Della Libera, Plomp, & Chelazzi, 2019; Suárez-Pinilla, Seth, & Roseboom, 2018). SD has also been described as a perceptual effect, with stimuli truly appearing to observers as more like previous stimuli (Cicchini, Mikellidou, & Burr, 2017; Cicchini, Benedetto, & Burr, 2021; Collins, 2019, 2020; Fritsche, Spaak, & de Lange, 2020; Kim, Burr, Cicchini, & Alais, 2020; Manassi, Kristjánsson, & Whitney, 2019; van Bergen & Jehee, 2019). Finally, SD has also been characterized as a memory effect resulting from the re-activation of previous stimulus representations during the processing of current stimuli and response preparation (Bae and Luck, 2019; Barbosa, Stein, Martinez, Galan-Gadea, Li, Damau, Adam, Valls-Solé, Constantinidis, & Compte, 2020; Bilacchi, Sirius, Cravo, & de Azevedo, 2022; Bliss, Sun, & D’Esposito, 2017; de Azevedo Neto and Bartels, 2021; Fritsche, Mostert, & de Lange, 2017; Mei, Chen, & Dong, 2019; Papadimitriou, Ferdoash, & Snyder 2015; Sheehan and Serences, 2022; Stein, Barbosa, Rosa-Justicia, Prades, Morató, Galan-Gadea, Ariño, Martinez-Hernandez, Castro-Fornieles, Dalmau, & Compte, 2020).

Similarly, repulsion has been described in several different contexts. For example, negative aftereffects are when prolonged exposure to a stimulus repulses the immediately following percept (Cicchini et al., 2021; Fritsche et al., 2017; Sadil, Cowell, & Huber, 2024). Repulsion can also manifest as a long-term effect, with stimuli in the distant past repulsing responses to the current stimulus (Chopin and Mamassian, 2012; Fritsche et al., 2020; Gekas, McDermott, & Mamassian, 2019; Suárez-Pinilla et al., 2018). Repulsion has also been reported in SD studies, previous stimuli far in feature space repulsing the perception of current stimuli (Bliss et al., 2017; Fritsche et al., 2017, 2020; van Bergen and Jehee, 2019; Samaha, Switzky, & Postle, 2019).

In all of the studies examining history effects, a sequence of stimuli with different spatial and temporal characteristics is shown to participants. Often, the designs causing repulsion and those causing attraction to appear highly similar. These contextual similarities in the appearance of repulsive versus attractive history effects raise the question of whether these effects are caused by the same underlying construct, or whether they are the expression of different mechanisms. One approach to try to reconcile the seemingly contrasting history effects in similar paradigms has been to consider attraction and repulsion as two sides of the same coin, suppressing one another (Fritsche et al., 2020) or co-existing (Alais, Leung, & Van de Burg, 2017; Taubert, Alais, & Burr, 2016; Pascucci et al., 2019). We set out to uncover if and how systematic differences in experimental design could lead to attraction or repulsion. We aimed to evoke serial dependence-like attraction (Fischer & Whitney, 2014) and negative aftereffect-like repulsion (Fritsche et al., 2017) in the same experiment by manipulating several variables.

The first variable of interest was the number of stimuli and responses in a trial. In studies on adaptation, a trial often has two stimuli, an adapter and a test. After an exposure to the adapter, subjects respond to the test, and the effect of the adapter on the test stimulus is measured. The effect is often repulsive, which is generally explained as the result of fatigue in adapter-sensitive channels and disinhibition of the surrounding channels, causing a net repulsion in the population-level response (for a review, see Webster, 2015). In contrast to this adapter-test design, most studies on serial dependence require subjects to respond to every stimulus. We propose that such a task requirement may promote (erroneous) visual continuity between sequential stimuli, resulting in attraction (Fischer and Whitney, 2014); while on the contrary, the adapter-test design may promote feature-differentiating processes and repulsion (Blondé, Kristjánsson, & Pascucci, 2023; Kiyonaga, Scimeca, Bliss, & Whitney, 2017). We hypothesized that responding to every stimulus would lead to attraction, and responding to every other stimulus (an adapter-test design) would lead to repulsion (see Czoschke, Fischer, Beitner, Kaiser, & Bledowski, 2018).

The second variable of interest was stimulus duration. The adaptation literature (for a review, see Kohn, 2007) suggests that repulsion is stronger when the adapting stimulus is longer than the test stimulus (Gibson and Radner, 1937; Harris and Calvert, 1989; Kanai and Verstraten, 2005; see also Experiment 7 in Fischer and Whitney, 2014). In the serial dependence literature, stimuli often have only one duration. Indeed, attraction may prevail in this case, because the design mimics the temporal regularities of a stable environment (Blondé et al., 2023; Gallagher and Benton, 2020; van Bergen and Jehee, 2019). We hypothesized that repulsion would emerge when the first stimulus was longer than the second, and attraction would emerge when stimuli had the same duration.

The third variable of interest was response type, by which we mean the way subjects reported their percept. In many serial dependence studies, subjects give a continuous adjustment response, and in others, a discrete categorical response. We were interested in examining to what extent these response types could influence the size and direction of the history effects. Some have examined this issue before. For example, Cicchini, Mikellidou, and Burr (2018) asked the same subjects to give a two-alternative forced choice (2AFC) and an adjustment response, but comparisons between the tasks are complicated by a response delay for the 2AFC task that was not present for the adjustment task. Fritsche et al. (2017) also gathered different response types: subjects adjusted a continuous response cue to match the orientation of a Gabor patch and then gave a 2AFC response comparing two Gabor patches. When subjects gave an adjustment response, their reports were attracted to the previous Gabor. When they had to compare orientations between two Gabors, one of which had been preceded at the same location by a cued Gabor, the previous cued Gabor worked as an adaptor and caused the Gabor in the 2AFC task to be repulsed away from the comparison Gabor. Fritsche et al. (2017) argued that the attractive effect with the adjustment response was post-perceptual because it was not location-specific (see Collins, 2019 showing the spatial extent of attraction). Conversely, the repulsive effect with the 2AFC response was argued to be perceptual because it resulted from comparing two stimuli and was location-specific. However, their design confounded response type (i.e., adjustment and 2AFC) and response order (i.e., first response adjustment and second response 2AFC). It may equally be likely that response type caused the effects or response order, with subjects needing to differentiate the stimuli in the 2AFC task from a previous stimulus in the adjustment task.

Forced choice tasks are sometimes seen as preferable to adjustment tasks for their direct approach (Pascucci et al., 2019; Pascucci et al., 2023; but see Yildirim, Coates, & Sayim, 2020 for a different opinion), but they suffer from their own potential biases, such as choice biases (Abrahamyan, Silva, Dakin, Carandini, & Gardner, 2016). To capture the history effects that cannot be attributed to a specific measure and unconfound response type and order (e.g., Fritsche et al., 2017), we asked our subjects to give categorical (i.e., forced-choice) and continuous (i.e., adjustment) responses in similar but separate tasks to gain insight into whether response type plays a systematic role. We did not have any strong hypotheses about whether one or the other response type would preferentially lead to repulsion or adaptation.

The fourth and final variable of interest was the origin of history effects. Several studies showed distinct effects from the previous stimulus and the previous response (Bosch et al., 2022; Pascucci et al., 2019), and we thus analyzed both effects as well. In this way, we hoped to establish the design variables that determine distinct history effects but also explore how they may co-occur.

To summarize, we manipulated experimental variables that could cause the repulsion or attraction in the perception (response) of (to) oriented Gabors: response type (2AFC and adjustment), response frequency (every trial and every other trial), and stimulus duration (50 and 500 ms). Our hypotheses were the following: (1) There would be repulsion with alternating responses (because such responses encourage subjects to differentiate the two stimuli), and attraction with sequential responses (because such sequential stimuli can be seen as continuous and promote integration); (2) repulsion would be stronger with longer previous stimulus duration and shorter current stimulus duration (inspired by the adaptation literature), and attraction would be stronger when both previous and current stimuli had the same stimulus durations (inspired by the serial dependence literature). We did not have a specific hypothesis regarding the response types but were simply interested in exploring how they may interact with other variables. In analyzing the data, we examined whether history effects differed when considering the influence of the previous response and the previous stimulus on the current trial.

## Methods

### Subjects

Fifteen subjects (10 female) aged 18 to 40 years (M=27.7, SD=6.2) with normal or corrected-to-normal vision participated in the experiment. All provided written informed consent prior to their participation and were compensated for their time (10€ per hour). The study was approved by the French ethics committee (Comité de Protection des Personnes Ouest IV-Nantes) and followed the ethical guidelines of the Declaration of Helsinki (2013).

### Stimuli and apparatus

Subjects sat in a dark room at a distance of 57 cm from a 328×556 mm, 1920×1080 pixel resolution, 240 Hz refresh rate monitor. Responses were collected using a keyboard and a mouse placed at a comfortable distance by subjects.

The stimuli were generated and presented in MATLAB R2021b using the Psychophysics Toolbox extensions (Brainard, 1997; Kleiner, Brainard, & Pelli, 2007; Pelli, 1997). All stimuli were presented at screen center on a noise background consisting of two-pixel squares in two shades of gray randomly selected. A 0.37 degree of visual angle (dva) diameter black fixation dot appeared at screen center. Gabor patches were presented centrally with 100% Michelson contrast, 1:1 aspect ratio, and 1.65 cycles per degree of spatial frequency and were windowed in a 0.76° standard deviation Gaussian envelope. Gabors had the same parameters in all experiments except for their orientation, which varied between ±30° and ±90° relative to the vertical 0°. Each subject was randomly assigned to either a clockwise (+) or counterclockwise (-) orientation distribution before their participation, which remained fixed in all sessions. A Gabor oriented ±60° depending on the direction of orientation distribution was the reference stimulus in the categorical response task.

There were three types of response cues. In the continuous response task, the response cue consisted of two white 0.33 dva-diameter dots placed symmetrically on an invisible 1.97 dva-diameter circle. To prevent the response cue from giving orientation information that could influence the remembered orientation of the Gabor, the initial positions of the dots were at screen center and expanded out to the invisible circle as subjects moved the mouse cursor. In the categorical response task, the change in the pattern of noise background following the stimulus presentation cued for a response. In the control trials (see <u>Procedure</u>), the response cue was a stationary red 0.36 dva-diameter dot at screen center.

### Procedure

Each subject participated in four sessions for each combination of response type and response frequency. Response type refers to the categorical response task and continuous response task. In the categorical response task, subjects reported whether the Gabor was clockwise (CW) or counter-clockwise (CCW) relative to a ±60° reference stimulus by pressing on the left or right arrow keys. In the continuous response task, subjects adjusted a response cue to match its orientation to that of the Gabor and registered their response with the left mouse click. Response frequency refers to whether subjects were required to give a sequential response (i.e., respond to every Gabor) or an alternating response (i.e., respond to every other Gabor). In the alternating response blocks, control trials ensured that subjects attended to the first Gabor by asking them every 5 to 10 trials to judge with a key response whether the second Gabor was oriented CW or CCW relative to the first (instead of the reference).

The sessions were categorized as continuous-sequential, continuous-alternating, categorical-sequential, and categorical-alternating. Their order was counterbalanced across subjects. Each lasted between 40 and 90 minutes and could be separated by a few days, and each consisted of 20 blocks of 61 trials (or 67 if control trials were present), totaling 5120 trials per subject across all sessions.

A trial started with the fixation dot on a noise background presented for 750 ms (see <u>Figure 1</u>). Then, an oriented Gabor randomly selected from a uniform distribution appeared for either 50 or 500 ms. At Gabor offset, the noise background changed to a new pattern. It thus served as a mask to prevent afterimages and a cue to respond. With alternating responses, after an inter-stimulus interval (ISI) of 500 ms, another Gabor appeared for 50 or 500 ms before the noise background changed, and Gabor offset cued to respond. The next trial started with the appearance of the fixation dot immediately after the response was given. A trial took 1350 or 2250 ms with alternating responses and 800 or 1250 ms with sequential responses (excluding response time). Subjects practiced the tasks at the start of each session in a block of 10 trials with sequential responses and 20 trials with alternating responses. If they failed to understand the tasks, subjects could request an additional practice block. This happened only twice (for Subject 2 in the categorical response task with alternating responses and Subject 14 in the continuous response task with sequential responses). In these trials, subjects were given feedback on their responses as follows. In the continuous response task, a red wedge appeared showing how far their report was from the actual orientation. In the categorical response task, a green or red text appeared for correct and incorrect responses, respectively. In the case of an incorrect response, the reference stimulus was presented again. During the experimental blocks, the reference stimulus was presented every 21st trial and between the blocks to prevent changes in the remembered orientation of the reference stimulus. Subjects determined the duration of these reference presentations with a key press to continue the experiment.

**Figure 1.**
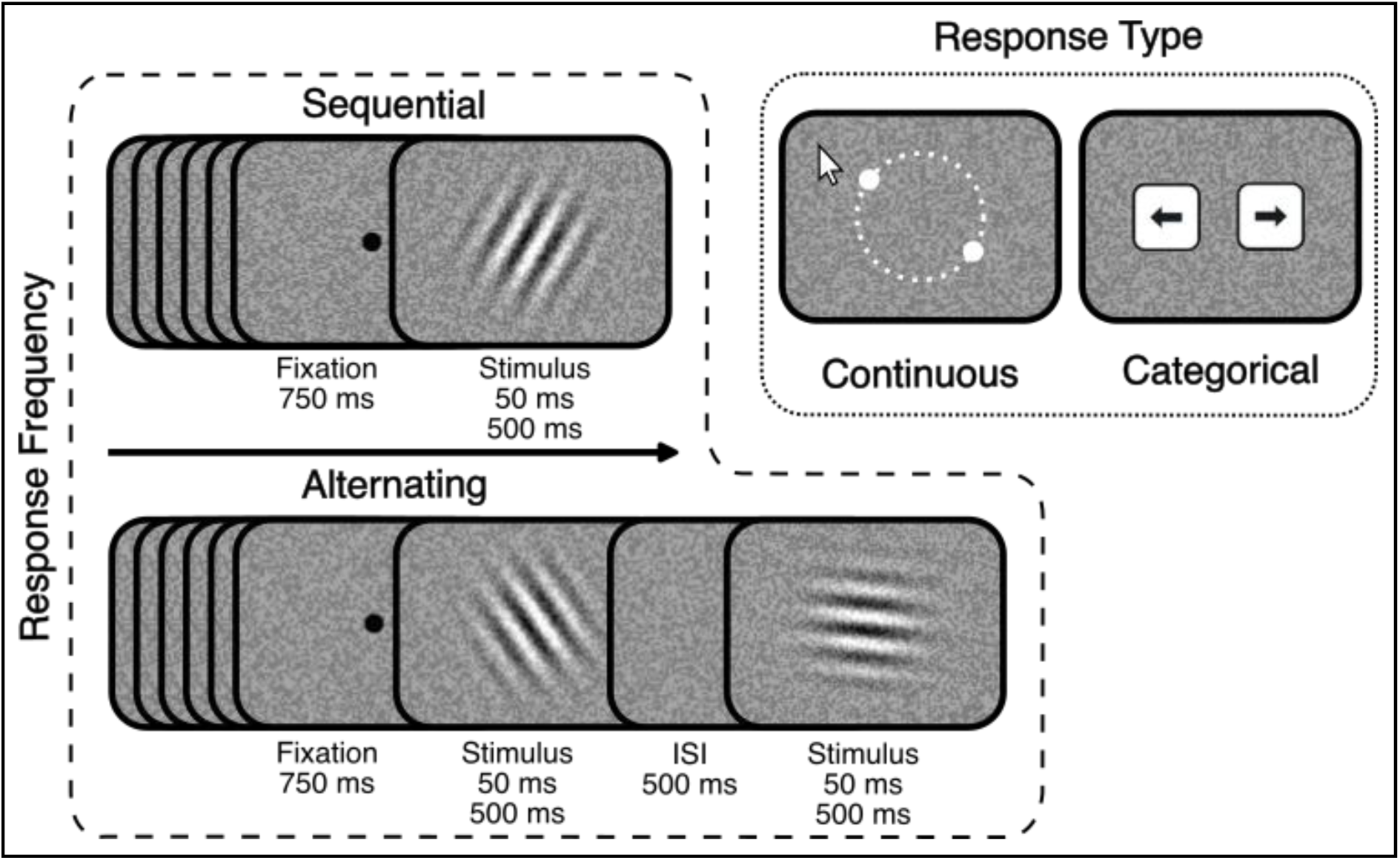
Experimental procedure. Stimuli sizes are not scaled. After a 750 ms fixation dot, an oriented Gabor was presented for 50 or 500 ms. With sequential responses, the response display appeared immediately with a new noise background. With alternating responses, a 500 ms interstimulus interval (ISI) followed the first Gabor and changed the noise background, after which another oriented Gabor appeared for 50 or 500 ms, and at its offset, the response display appeared with a new noise background. Subjects were instructed to respond either by reporting with a keypress whether the (second) stimulus was clockwise or counter-clockwise relative to a reference in the categorical response task or by adjusting a response cue with the mouse to match the (second) stimulus orientation in the continuous response task.

<u>Figure 1</u> shows all tasks and conditions. There were four independent variables: response type (continuous and categorical), response frequency (sequential and alternating), stimulus duration (50 and 500 ms), and stimulus orientation (±30° to ±90°). Response type and frequency varied between sessions, stimulus duration varied between blocks, and stimulus orientation varied between trials.

### Data analyses

We calculated CW response proportions in the categorical response task (i.e., total number of CW responses divided by total number of responses) and response errors in the continuous response task (i.e., orientation difference between the response and the stimulus). CW and CCW errors correspond to positive and negative values, respectively.

To find the influence of the past, we investigated trial sequences that were informative about the previous and current stimuli and responses. This meant that some data needed to be removed by default, such as the first trials of blocks because they did not have a relative orientation of previous stimulus and response and trials showing the same stimulus as in the previous trial. Additionally in the categorical task, trials after the reference display (i.e., every 21st trial of a block) were removed because they disrupted the sequence. Trials showing the same stimulus as the reference in the categorical response tasks were also removed for simplicity in the analysis (see the next paragraph describing the current stimulus orientation levels). Moreover, we excluded the trials following a control trial in tasks with alternating responses because the previous response in a control trial did not match the response in the current trial due to differences in task instructions (i.e., report the second stimulus vs. report whether the second stimulus is CW or CCW compared to the reference). Lastly, we removed trials if subjects did not move the cursor at all in the continuous response task and instead registered invalid responses. Then, we detected and removed outliers in responses using the jack-knife technique: After grouping for each subject, means and standard deviations were calculated with a leave-one-trial-out procedure, and if the left-out trial was more than ±3 standard deviations from the mean, it was excluded. This technique was applied to response times in the categorical and continuous response tasks and additionally to response errors in the continuous response task. If a trial was excluded, the following trial was also excluded because we wanted to exclude any uncontrolled effect exerted by an outlier trial. The jack-knife technique removed 3% of the trials in the categorical response task with alternating responses, 3.5% of the trials in the categorical response task with sequential responses, 4.2% in the continuous response task with alternating responses, and 3.7% in the continuous response task with sequential responses, leaving with 15328 (1095±21), 17112 (1101±11), 14278 (1020±18), 17982 (1154±16) trials (per subject), respectively. The reason for the large differences in the total number of trials between the tasks was that one subject failed to complete the tasks with alternating responses and had data only for sequential responses.

To test for statistical significance, we used linear mixed-effects models. We opted out of the canonical serial dependence analysis with derivatives of Gaussian models because the data included a limited range of relative orientations of previous stimulus (−60° to 60°), which fell on the slope of a first derivative of Gaussian function without providing data to model the extremities. This can be seen in <u>Figure 2</u> plotting the residualized response error used in the analysis (see also <u>Removing biases</u> in the Results section below for the residualization procedure). The linear mixed-effects models were built using R Statistical Software, in particular the *lme* function from the *nlme* package (Pinheiro, Bates, & R Core Team, 2024). Specific models are detailed in the appropriate part of the results section, which included some or all of the following fixed-effect factors: response type (continuous and categorical), response frequency (respond to every stimulus and respond to every other stimulus), previous stimulus duration (50 and 500 ms), current stimulus duration (50 and 500 ms), current stimulus orientation category (CW and CCW), relative orientation category of previous stimulus (CW and CCW), relative orientation category of previous response (CW and CCW), previous response (CW and CCW), and history effect origin (previous stimulus and response). We estimated degrees of freedom and computed p-values for the fixed effects in these models with Kenward-Roger approximation using the *model_parameters* function from the *parameters* package (Lüdecke, Ben-Shachar, Indrajeet, & Makowski, 2020). Significant interactions were followed up by pairwise comparisons with Bonferroni correction. For all comparisons, the alpha value was set to 0.05.

**Figure 2.**
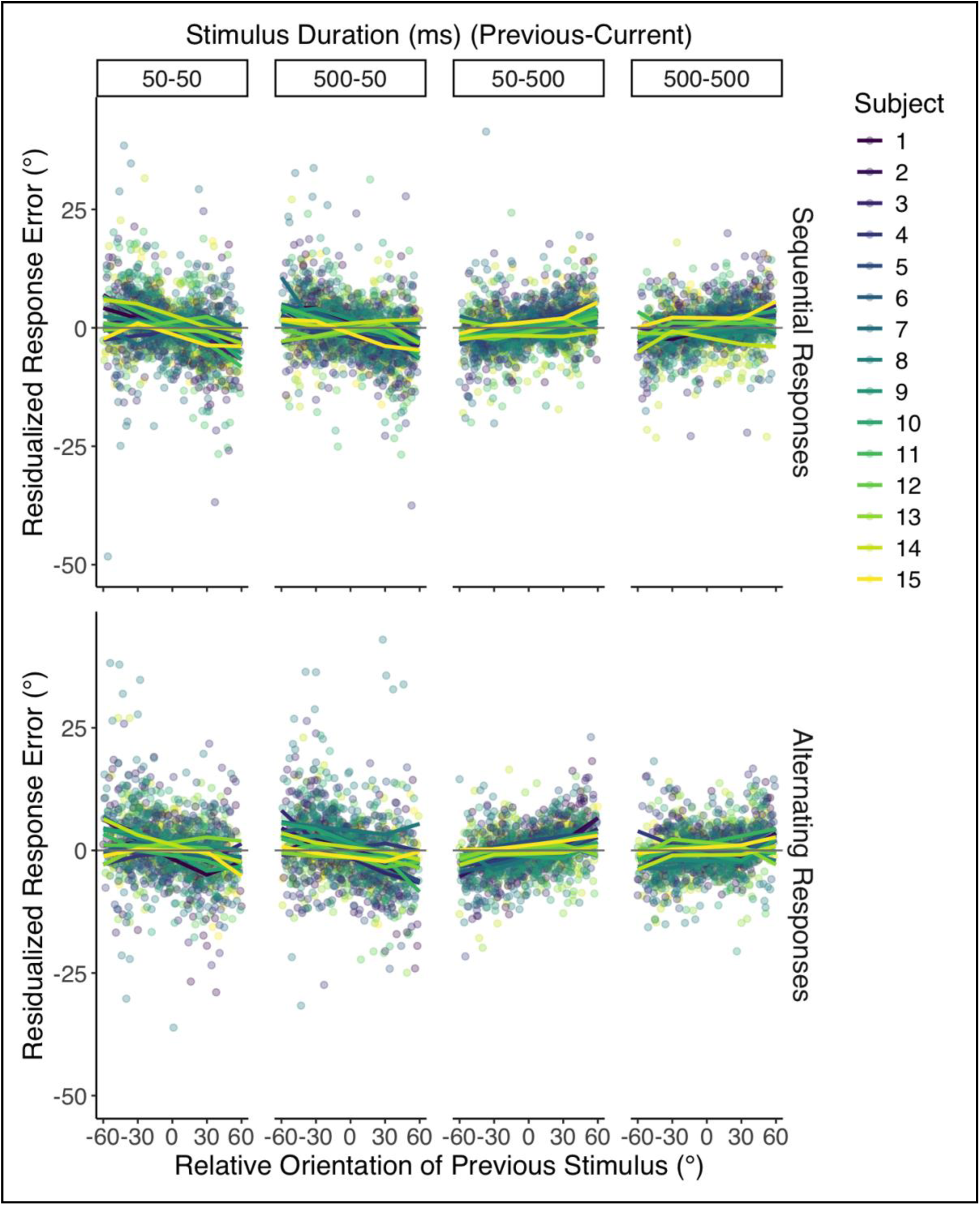
Continuous response task. Residualized response errors as a function of relative orientation of previous stimulus, response frequency, and stimulus durations of previous and current stimuli. The points show the averaged values for all subjects and relative orientations of previous stimulus, and the lines show the running averages of relative orientations in windows of 30°. Colors represent individual subjects.

### Data availability

The MATLAB scripts for data collection, the R scripts for data analysis, and the data will be made freely available upon acceptance. The design and analysis were not pre-registered on a public repository.

## Results

### Task performance

Response error magnitude in the continuous response task (i.e., the absolute mean difference between reported and actual orientations) and error proportion in the categorical response task (i.e., number of incorrect responses divided by total number of responses) are shown in <u>Figure 3</u>. Low error magnitudes and proportions show that subjects were able to perform the tasks relatively well. In the continuous response task, error magnitudes were a little larger for 50 ms (8°±1.8) compared to 500 ms (6°±1.4). This means that the task became slightly more difficult, as expected, when stimuli were presented briefly.

**Figure 3.**
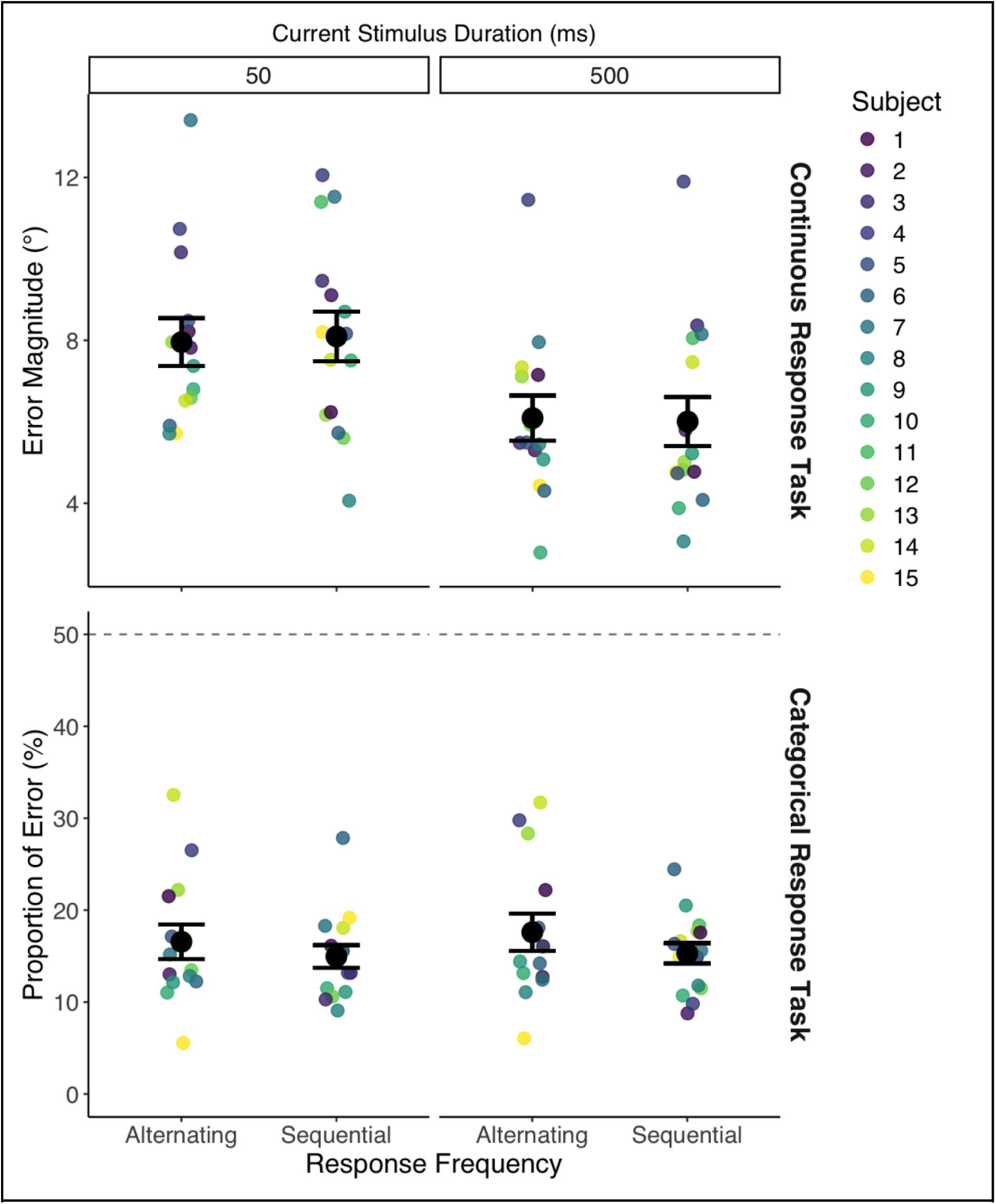
Response error magnitude in the continuous response task (top panel) and response error proportion in the categorical response task (bottom panel) as a function of response frequency and current stimulus duration. Colored points show subject averages across all trials. Larger black points show the average across all subjects. Error bars are standard errors. The dashed line in the bottom panel shows chance level performance.

Errors, as a proxy of task difficulty, did not show any other systematic differences between conditions. Thus, the effects reported in the following sections cannot be attributed to differences in task difficulty.

Accuracy in the control trials, which were only present in alternating response blocks, was high (74±3% across all subjects and conditions), suggesting that subjects were able to judge whether the current was CW or CCW relative to the previous and did not simply ignore the previous stimulus.

### Removing biases

In the continuous response task, we expected to observe a cardinal bias (i.e., responses away from the horizontal orientation, 90°), an oblique response bias (i.e., responses towards the oblique orientation, 45°), and a central tendency bias (i.e., responses away from the two ends of the distribution, 30° and 90°, towards the central orientation, 60°). These biases may confound the effects of the previous stimulus. Thus, we fit third-degree polynomials for each subject and session, and residuals were used in the analyses. This analysis step was further motivated by the observation that, after shuffling the order of trials and permuting a new temporal order 1000 times (Pascucci et al., 2019), which would abolish the effects of trial history, we observed central tendency and cardinal biases. These biases disappeared when residualized response error was plotted as a function of the current stimulus orientation. For the remainder of the manuscript, response error refers to the residualized response error.

### Continuous response task: Current stimulus duration determines whether the past exerts a repulsive or attractive effect

<u>Figure 4</u> shows CCW (negative) and CW (positive) response errors in the continuous response task. CW errors when the *previous stimulus and response* are CCW relative to the current stimulus (dotted green bars) indicate repulsion. This can also be seen with CCW errors when the previous stimulus and response are more CW (solid blue bars). Such repulsive patterns can be seen when the current stimulus duration was 50 ms across response frequencies and previous stimulus durations. With the longer stimulus duration (500 ms), repulsive effect switched to attraction. CW errors when the *previous stimulus and response* were more CW and CCW errors when the *previous stimulus and response* are more CCW attest this. Such attractive patterns also hold across response frequencies and previous stimulus durations. Contrasting the opposing orientations of previous stimuli and responses, we can see their isolated effects (in solid green bars and dotted blue bars). This shows us that the attraction to the previous response works against the attraction to the previous stimulus when the current stimulus duration was long (thus values closer to zero).

**Figure 4.**
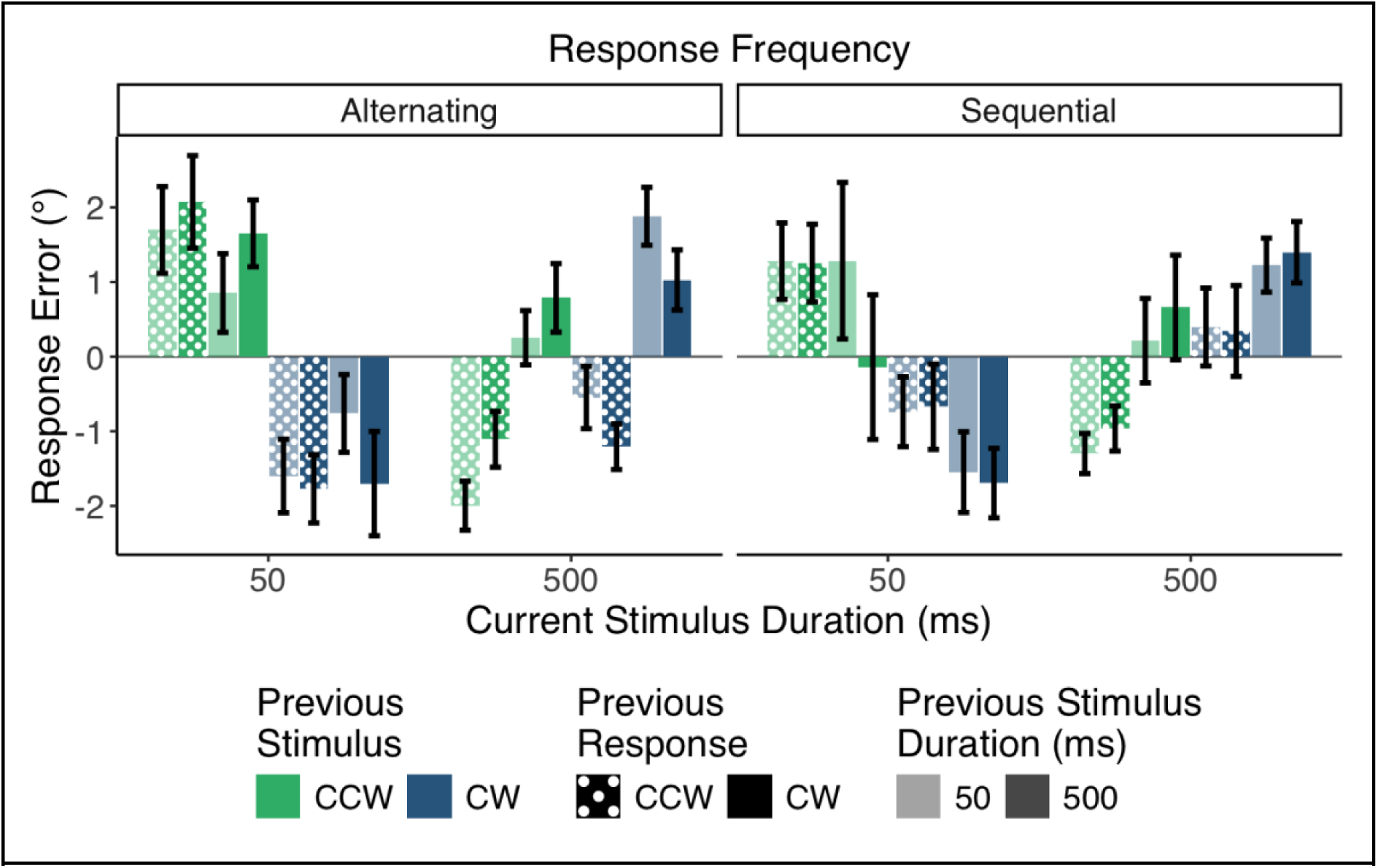
Averaged response errors of the continuous response task as a function of previous response (in patterns), previous stimulus (in colors), previous stimulus duration (in shades), current stimulus duration (in x-axis), and response frequency (in columns). Error bars show standard errors.

We confirmed these patterns with a linear mixed-effects model. First, we labelled the relative orientation of previous response (i.e., the distance between previous response and current stimulus orientation) and the relative orientation of previous stimulus (i.e., the distance between previous and current stimuli orientations) as CCW (−) or CW (+). These variables will respectively be referred to as *previous response* and *previous stimulus* for the remainder of the manuscript. Then, we included them in the model as fixed effects along with response frequency (sequential and alternating), previous stimulus duration (50 and 500 ms), and current stimulus duration (50 and 500 ms); subject as a random-intercept effect. Response errors were averaged across all levels of each term in the model, and the model was fit as follows:

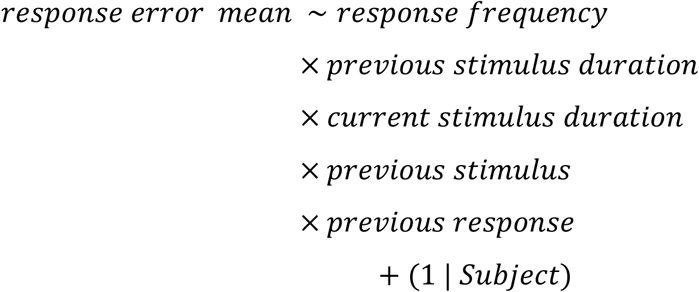

The linear mixed-effects model revealed a significant interaction between the current stimulus duration and the previous stimulus, *t*(417) = 4.36, *p* = 0.001 x 10^-2^, and between the current stimulus duration and the previous response, *t*(417) = 2.85, *p* = 0.005. Pairwise comparisons between CCW versus CW previous stimuli and responses for each current stimulus duration and between current stimulus durations for each previous stimulus and response showed significance (all *p* < .001) except for the comparison among previous responses when the current stimulus was presented for 50 ms (for which *p* = 0.57).

<u>Figure 5A</u> summarizes the findings from the continuous response task by collapsing data across non-significant factors, keeping only current stimulus duration, previous stimulus and response. <u>Figure 5B</u> plots the difference in response errors between CCW and CW previous stimuli/responses in <u>Figure 5A</u>, which we call the cumulative response error. These figures confirm the analysis detailed above: when the current stimulus duration is short, there is only attraction to the previous stimulus; when the current stimulus duration is long, there is attraction to both previous stimulus and response.

**Figure 5.**
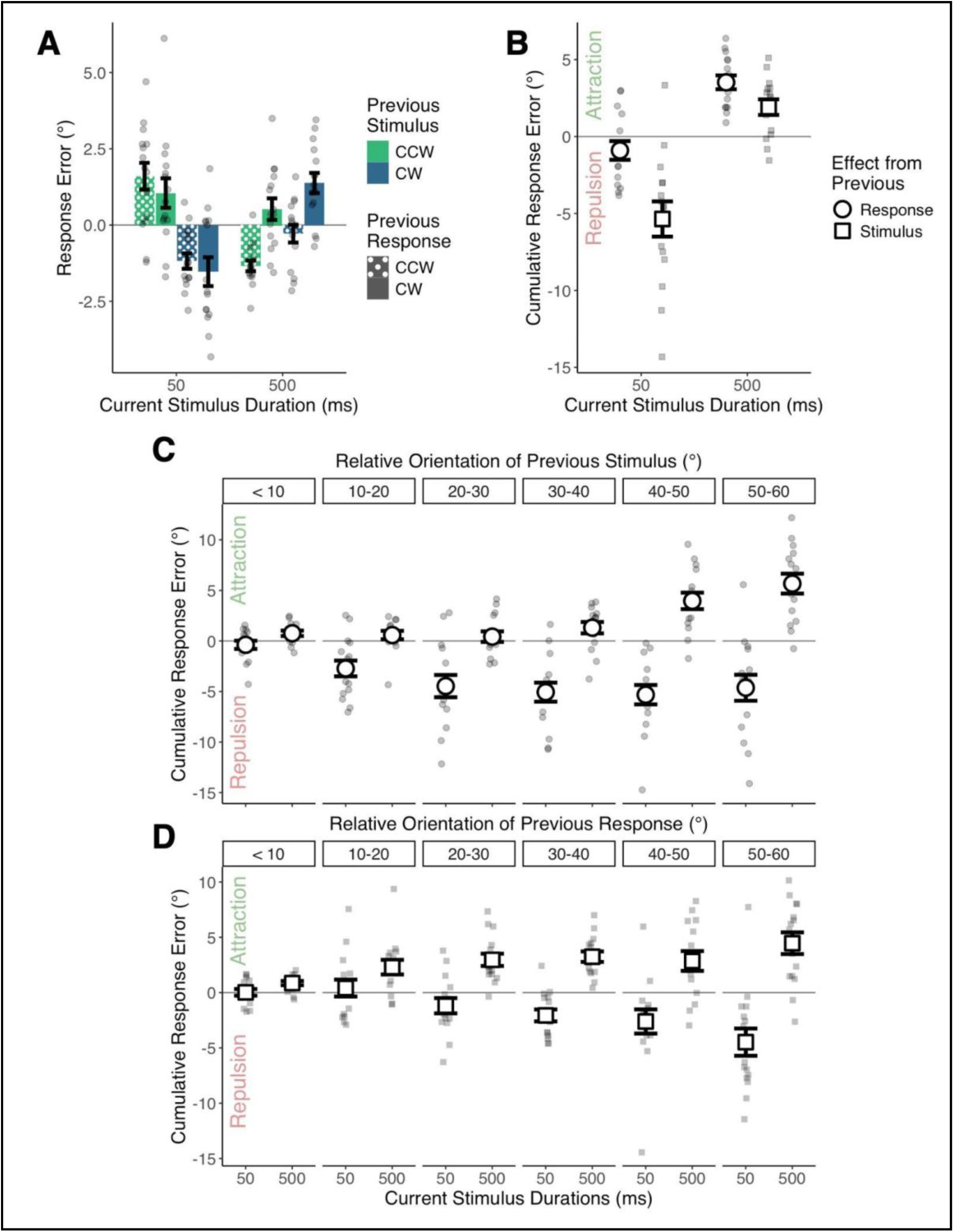
Continuous response task. (A) Average response error as a function of current stimulus duration, previous stimulus and response. (B) Cumulative response error across previous stimuli and responses as a function of current stimulus duration. (C) Cumulative response error of previous stimulus as a function of current stimulus duration and relative orientation of previous stimulus binned in degrees of 10. (D) Cumulative response error of previous response as a function of current stimulus duration and relative orientation of previous response binned in degrees of 10. Smaller points show averaged values for subjects, and error bars are for standard errors.

<u>Figure 5C-D</u> shows that the history effects (i.e., repulsion when the current stimulus lasted 50 ms and attraction when it lasted 500 ms) increased with the actual relative orientation between successive events.

These figures confirm the well-known feature tuning of serial dependence: the influence of the previous stimulus grows with distance from the current stimulus - to a certain threshold. We additionally show this feature when considering the distance between previous response and current stimulus. We do not observe a return to a response error of zero, probably because our design did not sample very large relative orientations. This feature tuning holds both when the history effect was attractive (long current stimuli durations) and when it was repulsive (short current stimuli durations).

These results reject our hypothesis that repulsion and attraction result from both current and previous stimulus duration (hypothesis #2, see <u>Introduction</u>) because only current stimulus duration determined which history effect manifested. They also reject our hypothesis (#1) that alternating and sequential tasks would lead to repulsion and attraction, respectively. Nevertheless, repulsion with stimuli whose duration is as short as 50 ms (Fornaciai & Park, 2019) and attraction with stimuli as long as 500 ms (Fischer & Whitney, 2014), both of which modulate as a function of orientation distance between the previous and current stimuli, are in line with the literature. We also show that even when responding only to every other stimulus, attraction to the past still occurs (Fischer and Whitney, 2014; Suárez-Pinilla et al., 2018). Unlike previous reports showing opposing effects of previous stimulus and response (Moon and Kwon, 2022; Pascucci et al., 2019; Sadil et al., 2024; Zhou, Liu, Jiang, Wang, Xu, & Zhou, 2024), however, we found an additional attraction to the previous response with the longer stimulus duration.

### Categorical response task: Previous responses cause attraction and previous stimuli repulsion

<u>Figure 6</u> shows the proportion of CW responses in the categorical response task. When the current stimulus is CW (blue and purple bars), we expect more CW responses than when the current stimulus is CCW (light and dark green bars), and this is evident in all conditions. Repulsion and attraction can be seen here as differences between conditions. The effect of the previous *stimulus* appears when contrasting CCW and CW previous stimuli: previous trials with a more CCW stimulus (light green and blue bars) have higher proportions of CW responses than previous trials with more CW stimulus (dark green and purple bars) – a repulsive effect. The effect of the previous *response* appears when contrasting previous CCW and CW responses: previous CW responses (all solid bars) have higher proportions of CW responses than previous CCW responses (all dotted bars) – an attractive effect.These effects were investigated by a linear mixed-effects model. Fixed effects were the previous response (whether it was CCW or CW), current stimulus orientation (CCW or CW relative to the reference), previous stimulus (CCW or CW – labels of the distance between previous stimulus and current stimulus orientation), current and previous stimulus durations (50 or 500 ms), and response frequency. The CW response proportions, averaged across the levels of each term in the model, were fit in the following fashion:

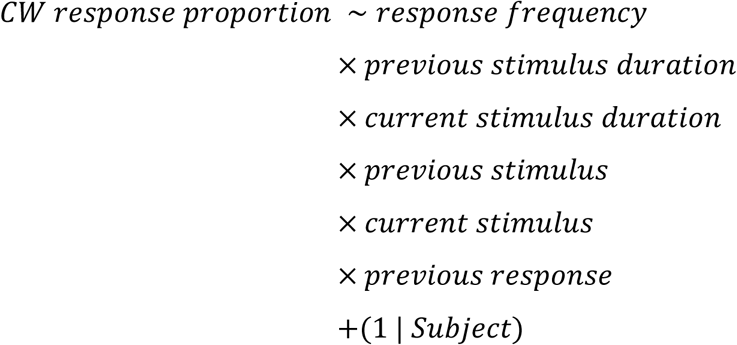

The main effect of the current stimulus was significant, *t*(823) = 9.79, *p* < 0.002 x 10^-13^, meaning that subjects were successful in performing the task. The main effect of the previous stimulus was significant, confirming the repulsive effect, *t*(823) = −2.67, *p* = 0.008. The previous response also reached significance, *t*(823) = 2.60, *p* = 0.01, confirming the attractive effect.

**Figure 6.**
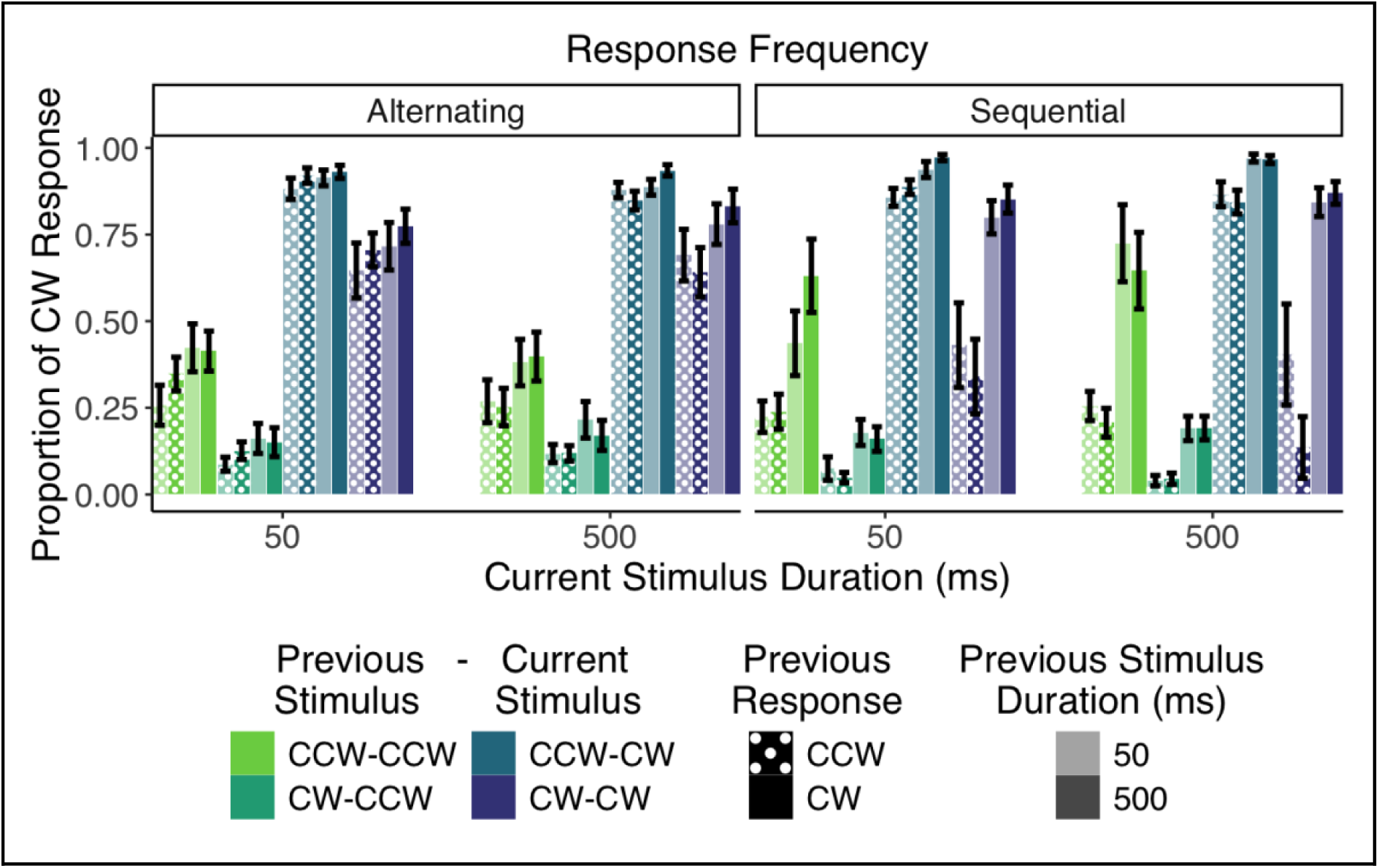
Averaged proportions of CW responses of the categorical response task as a function of previous response (in patterns), previous stimulus and current stimulus (in colors), previous stimulus duration (in shades), current stimulus duration (in x-axis), and response frequency (in columns). Error bars show standard errors.

<u>Figure 7A</u> summarizes the results for the categorical task by showing only the factors that significantly influence the proportion of CW responses (i.e., current stimulus, previous stimulus, and previous response). Here again, the repulsive effect of the previous stimulus is apparent: more CCW previous stimuli (light green and blue bars) lead to more CW responses than more CW previous stimuli (dark green and purple bars). The attractive effect of previous responses can be seen by comparing previous CW responses (solid bars) to the corresponding previous CCW responses (dotted bars): previous CW responses lead to more CW responses than previous CCW responses (significant pairwise comparisons between more CCW and CW of previous stimuli and between CCW and CW previous responses for each current stimulus, all *p* < 0.05). <u>Figure 7B</u> further synthesizes the results by subtracting the proportion of CW responses when the previous response (or stimulus) was CCW from when the previous response (or stimulus) was CW and summing them across both current stimulus orientations. This summarizes how previous stimuli repulse current responses, whereas previous responses attract them. <u>Figure 7C</u> further shows that the attraction from the previous response got weaker with increasing relative orientations while the repulsion from the previous stimulus increased.

**Figure 7.**
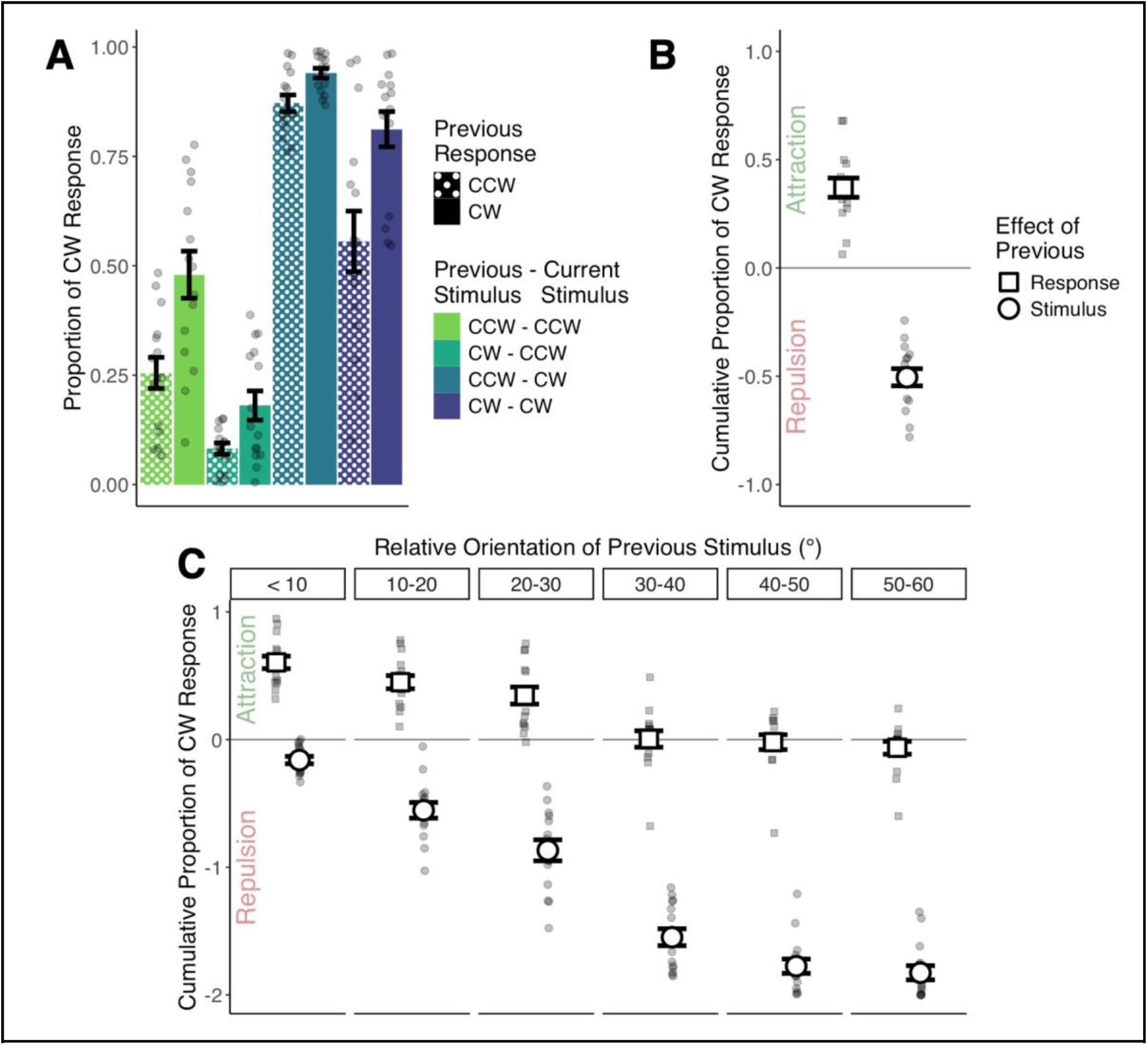
Categorical response task. (A) Proportions of CW response as a function of previous stimulus, current stimulus, and previous response. (B) Cumulative proportions of CW response (i.e., sum of differences between previous responses and previous stimuli across the current stimuli). (C) Cumulative proportions of CW response as a function of relative orientation of previous stimulus binned in degrees of 10. Smaller points show values for subjects, and error bars are for standard errors.

We further found that the attractive effect of the previous response was modulated by response frequency and stimulus durations as confirmed by three-way interactions between response frequency, previous stimulus duration, previous response, *t*(823) = 2.13, *p* = 0.034, and response frequency, current stimulus duration, previous response, *t*(823) = 2.19, *p* = 0.029. <u>Figure 8A</u> illustrates these effects by averaging out the other factors, including current stimuli. This results in proportions of CW responses around 50%, allowing us to see history effects as deviations from this mean. Significant pairwise comparisons between CCW and CW previous responses showed attraction to the previous stimulus across almost all of the conditions (all *p* < 0.05, except for alternating responses when current stimulus duration was 50, for which *p* = 0.13). There was a stronger attraction towards the previous response with sequential responses than with alternating responses, modulated by previous and current durations. This modulation is shown more neatly in <u>Figure 8B</u>, in which the proportion of CW responses when the previous response was CCW (dotted bars) was subtracted from when the previous response was CW (solid bars). Here, we can see that attraction was stronger with sequential responses, the longer current stimulus duration, and the longer previous stimulus duration. However, significance was reached only for the comparisons between response frequencies (all *p* < 0.05, except for when previous response was CW and previous and current stimulus durations were 50 ms, for which *p* = 0.1 and 0.51, respectively).

**Figure 8.**
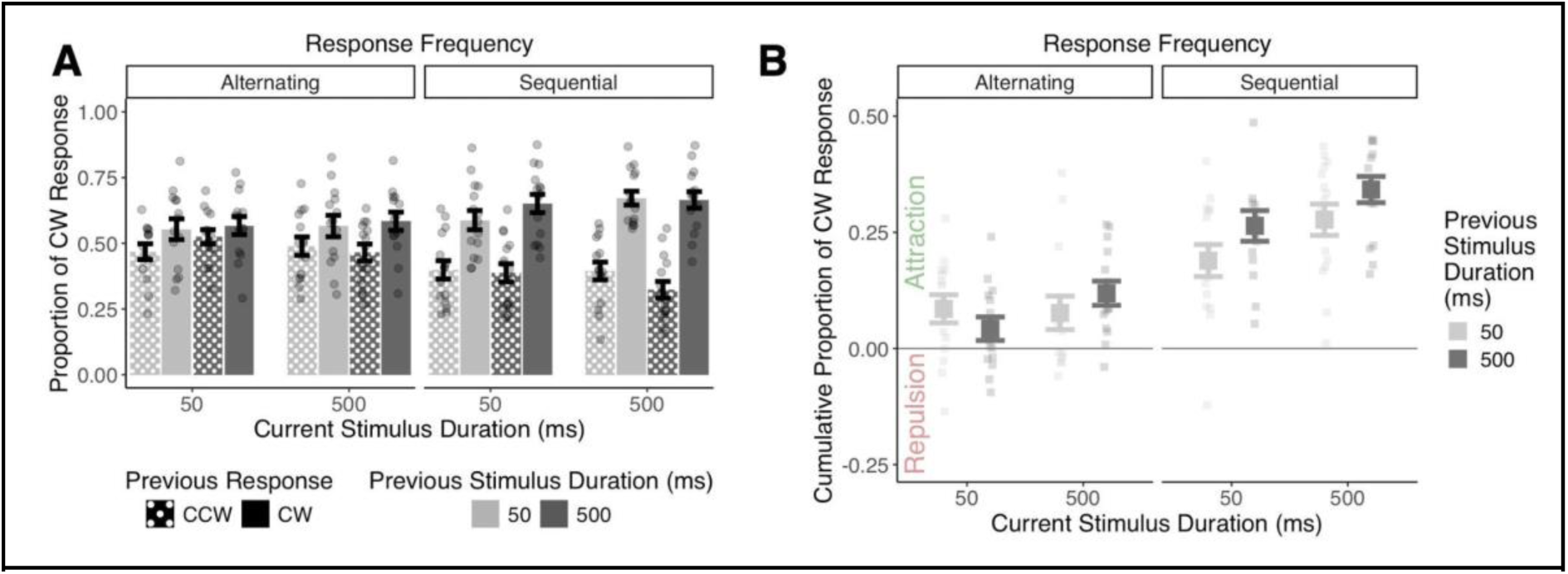
Categorical Response Task. (A) Proportion of CW response as a function of response frequency, previous stimulus duration, current stimulus duration, and previous response. (B) Cumulative proportion of CW response as a function of current and previous stimulus durations. Smaller points show values for subjects, and error bars are for standard errors.

These results reject our hypothesis (#1) that there would be attraction with sequential and repulsion with alternating responses. Hypothesis (#2) that the strength of effects would be modulated by the current and previous stimulus durations is also rejected. Nevertheless, we found opposing effects from the previous response and stimulus in a categorical task, complementing previous reports (Bosch, Fritsche, Ehinger, & de Lange, 2020; Bosch et al., 2022; Schwiedrzik et al., 2014; Van Geert et al., 2022). Moreover, attraction to the previous response was stronger with sequential responses compared to alternating responses, likely because the previous stimulus was closer in time with sequential responses, thus influencing the current response more strongly (akin to the inter-stimulus interval, see Bilacchi et al., 2022). This too shows that attraction to the past, limited to the response, still occurs when responding only to every other stimulus (Fischer and Whitney, 2014; Suárez-Pinilla et al., 2018), although to a lesser degree.

### Response type determines the variables facilitating attractive and repulsive effects

In the previous sections, we examined continuous and categorical response types separately. To explore how response types interacted with other variables, the cumulative effects reported in the previous sections were standardized by calculating the difference of each data point from the grand mean and dividing it by the standard deviation. <u>Figure 9</u> shows these standardized scores (i.e., z-scores), which quantify the history effect of the previous response and the previous stimulus across response types, stimulus durations, and response frequencies. We then ran a linear mixed-effects model on these z-scores to directly compare the response types (categorical and continuous). The history effects caused by the previous response and previous stimulus were labeled as levels of a new factor called “history effect origin” and added to the following model:

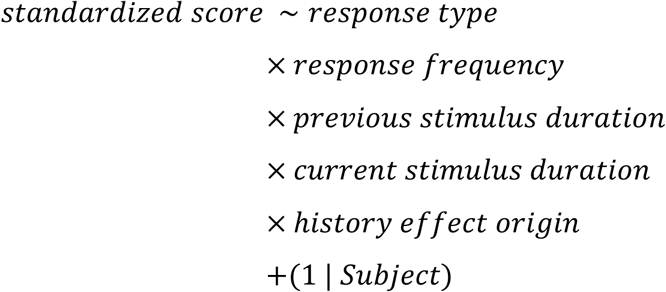

**Figure 9.**
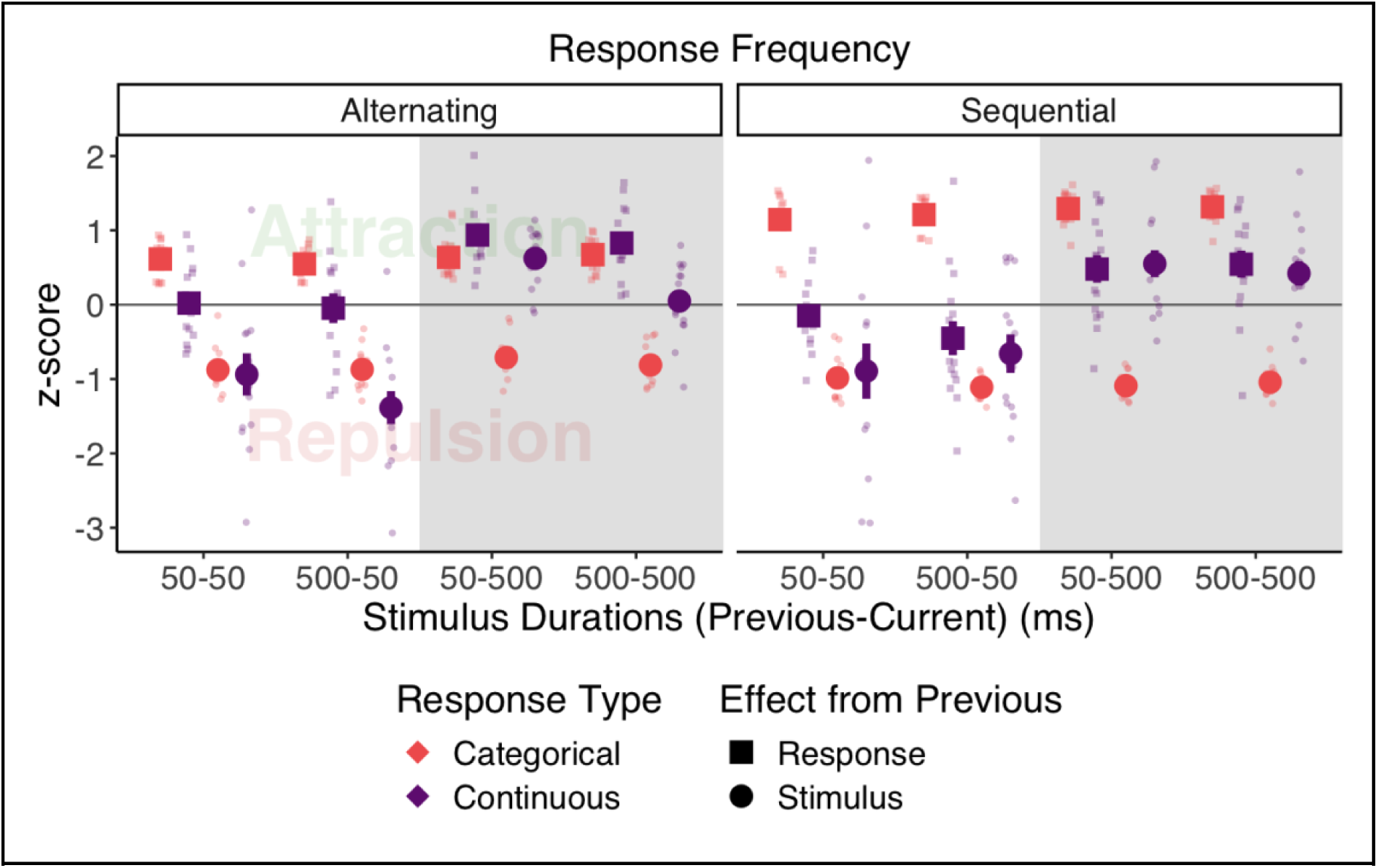
Standardized scores as a function of response type, response frequency, previous stimulus duration, current stimulus duration, and history effect origin. Positive and negative z-scores indicate attraction and repulsion, respectively. Smaller points represent subjects, and error bars are for standard errors (some occluded behind data points). The data points for the longer current stimulus duration (i.e., 500 ms) are highlighted in grey.

This analysis revealed the two ground rules: opposing effects with categorical responses (i.e., previous response led to repulsion and previous stimulus attraction), and additive effects with continuous responses (i.e., both previous response and stimulus led to the same effect but switched signs with the current stimulus duration). This was confirmed by a two-way interaction between response type and current stimulus duration, *t*(416) = 3.03, *p* = 0.003, showing that the repulsive effect from the previous trial switched to attraction for the longer current stimulus duration in the continuous but not in the categorical response task, because opposing effects canceled each other when averaging across history effect origins.

The model revealed several other interactions that highlight several modulations to these rules, as revealed by the four-way interaction between response type, response frequency, previous stimulus duration, and history effect origin, *t*(416) = 2.03, *p* = 0.043. First, previous responses led to attraction and previous stimuli repulsion across the response types, response frequencies, and previous stimulus durations (averaged values for squares in <u>Figure 9</u> are positive and for circles negative). Second, attraction to the previous response was stronger with categorical sequential responses (red squares were more positive in the right column), reiterating the response frequency effect reported for the categorical response task. Third, attraction to the previous response was absent with continuous sequential responses, because it was overridden by repulsion when the previous stimulus duration was long (purple squares in the right column are at zero when averaged across current stimulus durations). Similarly, repulsion from the previous stimulus in the continuous response task survived only with alternating responses when the previous stimulus duration was longer (purple circles in the left column are *not* zero for the 500 ms previous stimulus duration when averaged across current stimulus durations).

Several other lower-order interactions detail the same effects. For example, the model revealed a three-way interaction between response type, response frequency, and history effect origin, *t*(416) = 2.06, *p* = 0.04. This interaction comes from the lack of interaction between the response frequency and history effect origin when considering the continuous response task: history effects of response and stimulus did not change across the response frequencies. Meanwhile, there was an interaction between the response frequency and history effect origin when considering the categorical response task: previous responses caused stronger attraction with sequential responses, while repulsion from the previous stimuli remained the same across the response frequencies. A two-way interaction between response frequency and history effect origin, *t*(416) = −2.20, *p* = 0.028, also confirms this by showing that attraction to a previous response was better at overriding repulsion from a previous stimulus with sequential responses.

There was also a two-way interaction between response type and response frequency, *t*(416) = −2.41, *p* = 0.017. This interaction is illustrated in <u>Figure 10</u>. It reiterates the absence of an effect of response frequency in the continuous task. It additionally highlights that the combined effect of the previous categorical response and stimulus was repulsive with alternating responses and attractive with sequential responses (*p* = 0.043). Recall that in the categorical task, we observed attraction from the previous response and repulsion from the previous stimulus (see red data points in <u>Figure 9</u>). Here, when the two effects are combined, repulsion wins with alternating responses, and attraction wins with sequential responses. This suggests that the element - previous response or previous stimulus - closest in time to the current trial influences the current trial more strongly. In the case of the alternating task, the previous stimulus was closer in time than the previous response, thus the stimulus exerted its repulsive effect more. In the case of the sequential task, the previous response was closer in time than the previous stimulus, and it exerted its attractive effect the most.

**Figure 10.**
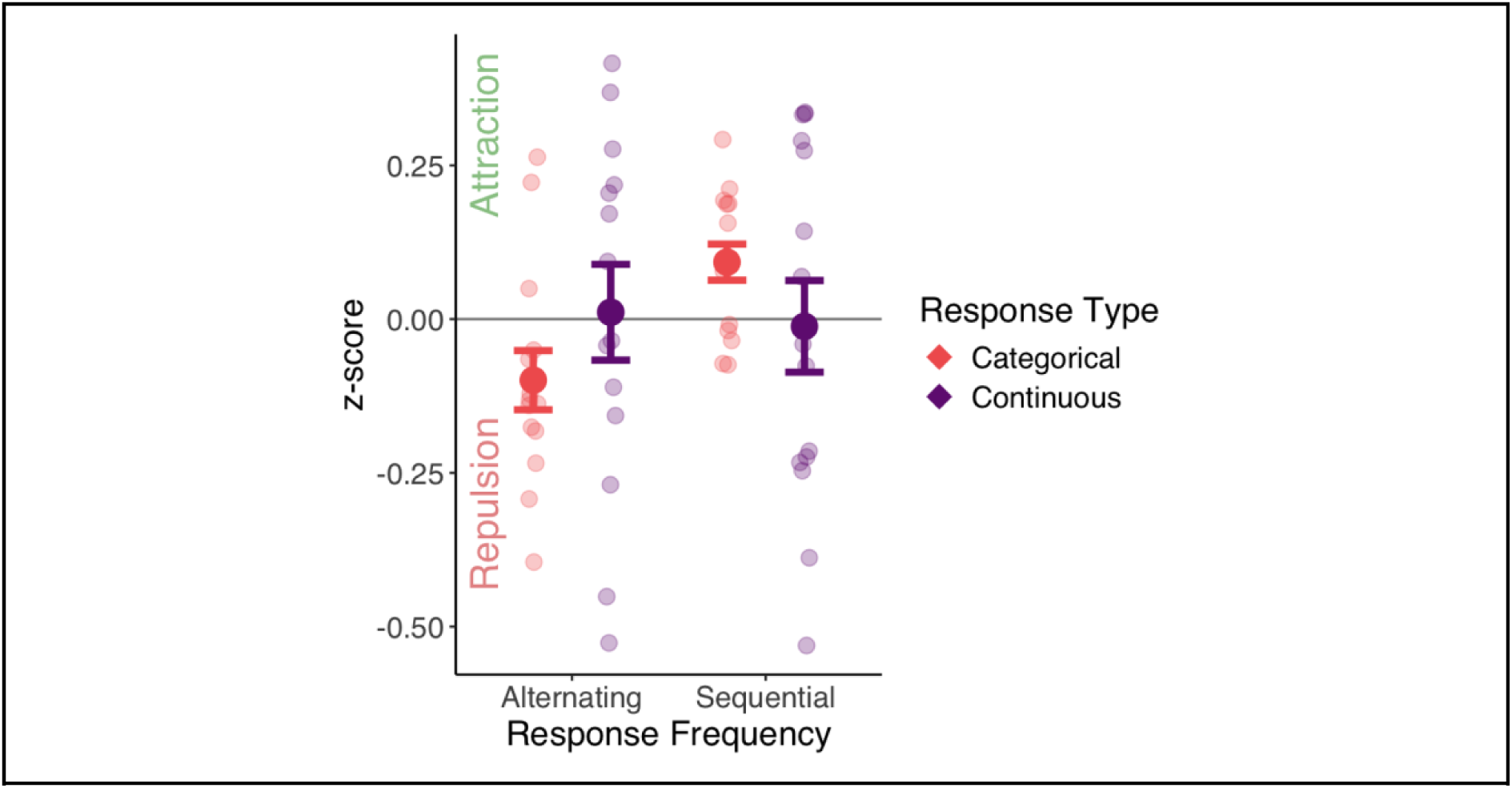
Standardized scores as a function of response type and response frequency. Smaller points show subject averages, and error bars are for standard errors.

These results show that response types interact with other variables and thus determine the variables facilitating either attractive or repulsive history effects. In the continuous response task, current stimulus duration determines the direction of the history effect of response and stimulus. In the categorical response task, response frequency modulates the strength of the history so much so that it renders the variable closest in time (previous stimulus or response) relevant in influencing the current response, which then determines the direction of the effect. Previous stimuli and responses can lead to attractive or repulsive history effects which may work together or in opposition. This suggests mechanisms that could switch flexibly between attraction and repulsion as a function of task demands.

## Discussion

We manipulated several experimental design variables to quantify their influence on the magnitude and direction of attractive (serial dependence) and repulsive (adaptation) effects with oriented Gabor patches. We varied response frequency (alternating and sequential) and stimulus duration (50 and 500 ms), making specific hypotheses about how these two variables would influence the strength and direction of history effects. We also varied response types (continuous and categorical), because they are both used in the serial dependence literature, to explore how they interact with other variables. Finally, we examined how both the previous response and stimulus influenced the responses to the current stimulus by including them as separate factors in our analysis.

The results rejected our main hypotheses. First, contrary to our expectation, we found that response frequency (i.e., responding to every stimulus versus responding to every other stimulus) was negligible in determining the *direction* of the history effects, but it influenced their *magnitude*. Second, again contrary to our hypothesis, we did not find that previous and current stimulus duration interacted to determine the direction of history effects. Instead, both previous and current stimulus durations independently interacted with response frequency in determining the strength of history effects. Third, we contrasted the influence of the previous response with that of the previous stimulus. Indeed, research has suggested that responses and stimuli may not exert the same effect on subsequent trials (Bosch et al., 2022; Pascucci et al., 2019). Our results confirm this, but only for the categorical response task. In contrast, we found that history effects of stimulus and response exert the same effect in the continuous response task and that the effect switches direction as a function of current stimulus duration. Lastly, we studied response types (categorical versus continuous) to measure if differences in task interacted with the other variables we manipulated. Not only did response type determine whether the previous response exerts the same effect as the previous stimulus or an opposing effect, but it also revealed the dynamics of history effects in interaction with the current stimulus duration, response frequency, and previous stimulus duration.

Why did we not find the response frequency effects we expected? We had hypothesized that responding to every other stimulus would lead to a repulsive effect because this design most resembles designs used in the adaptation literature (e.g., Kanai and Verstraten, 2005), in which one stimulus serves as an adapter and a second stimulus as a test to which subjects respond. One possibility is that our subjects simply did not pay attention to the first stimulus, since it did not require a response. We reject this possibility because we included control trials in which subjects had to compare the second stimulus to the first, requiring them to attend to the first stimulus. Performance in the control trials was high. Another possibility is that there was the expected repulsion but that it was overcome by the attraction. Indeed, when considering all variables in analyzing the response type effects, we observed an overall repulsion with alternating responses and attraction with sequential responses (see <u>Figure 10</u>). Variables that drive attraction with sequential responses were stronger than repulsion, and variables that drive repulsion with alternating responses were stronger than attraction.

Why did we not find an interaction effect of previous and current stimulus durations? We had hypothesized that the longer previous stimulus duration should lead to stronger repulsion, again because this design most resembles those used in the adaptation literature (Gibson and Radner, 1937; Harris and Calvert, 1989; Kanai and Verstraten, 2005), in which one (usually longer) stimulus serves as an adaptor and a second (usually shorter) stimulus as a test. We also hypothesized that when previous and current stimuli were of equal duration, we should observe stronger attraction, because this design resembles that used in the serial dependence literature (e.g., Fischer and Whitney, 2014) and it may contribute to a belief in continuity between trials (Blondé et al., 2023). As with the response frequency effect, stimulus durations alone failed to explain the strength of history effects. They instead independently interacted with the response type and frequency, showing a more complex pattern. There was stronger repulsion with the longer previous and shorter current stimulus durations, conforming with the adaptation literature, although this was limited to the continuous responses: stronger repulsion neutralized the attractive response effect with continuous sequential responses and surpassed the attractive stimulus effect with continuous alternating responses. However, we did not find stronger attraction with equal stimulus durations, suggesting that stimulus duration may play a negligible role in visual stability. Another explanation can be attributed to the block design (i.e., stimulus durations varied across the blocks). Subjects may have picked up on the regularity of changes in stimulus duration and assumed macro-level continuity instead of relying on micro-level changes in stimulus durations. This would explain why subjects had attraction at all stimulus durations except for conditions where repulsion could surpass it. Thus, our study is limited in answering what would have happened if stimulus durations were randomly varied within blocks.

Despite the lack of interaction between previous and current stimulus durations, our results reveal a crucial role of timing in determining the direction of history effects. First, the current stimulus duration strengthens attraction to the past. The fact that sequential responses to a series of successive 500-ms stimuli lead to attraction fits nicely with the large body of literature on serial dependence (e.g., Cicchini et al. 2017; Fischer and Whitney, 2014; Fritsche et al., 2017). Meanwhile, stimuli lasting for shorter durations were shown to be subject to repulsion from the past (Fornaciai and Park, 2019). This may resemble the response delay effect (Bliss et al., 2017; Fritsche et al., 2020). The more time there is for a past representation to influence the perception of and responses to the current stimulus, the stronger the attraction gets. Indeed, we show an additional attraction to the previous response appearing with the longer presentation times in the continuous response task. Second, response frequency determines the history effects as it is by definition confounded with time. Response frequency determined the event (previous stimulus or previous response) closest to the current trial, and perceptual reports about current stimuli were influenced by that recent event - previous response (attraction) or stimulus (repulsion). This may be likened to inter-stimulus interval effects (Bilacchi et al., 2022). The influence of a past representation decays with time. However, studies independently manipulating each of these variables (i.e., stimulus duration, response delay, response frequency, and inter-stimulus interval) are needed to fully determine their influence.

We also found that the previous response and stimulus independently contributed to history effects. The finding of attraction to the previous response co-existing with repulsion from the previous stimulus matches recent reports (Bosch et al., 2020; Bosch et al., 2022; Moon and Kwon, 2022; Pascucci et al., 2019; Sadil et al., 2024; Schwiedrzik et al., 2014; Van Geert et al., 2022; Zhou et al., 2024), although this was limited to categorical responses in our case. The previous stimulus and response exerted the same attractive effect on stimuli presented for longer with continuous responses. What causes this persistent reliance on the previous response across all conditions with categorical responses? One possible explanation could be decision confidence. Van Geert et al. (2022) found that subjects showed attraction toward responses to an ambiguous stimulus that lacked any stimulus information but to a lesser extent than responses to a stimulus with orientation information. They concluded that subjects’ responses were attracted towards more confident responses since their decisions were based on perceptual evidence. This agrees with a large body of studies (Bosch et al., 2020; Samaha et al., 2019; Suárez-Pinilla et al., 2018; see also Braun, Urai, & Donner, 2018). Thus, it may be that a continuous response measure is better at capturing what appeared to subjects thus contributing to the attractive effect of the previous stimulus and response, especially when they have more time to exert an effect. This would suggest that there may be distinct mechanisms at play from the previous response and stimulus and that other variables, determined by the response type, can influence these mechanisms.

The argument of two distinct mechanisms, one driven by the previous response and stimulus, was supported by the role of distance in feature space between two successive stimuli. The attraction to the previous categorical response increased with small relative orientations. This makes sense because as the feature distance grew, the probability that two successive trials were on two very different sides of the reference grew. In other words, subjects should give the same response twice in a row when the two successive stimuli are similar, independently of any history effects (i.e., choice repetition-like attraction). Meanwhile, the repulsion from previous categorical and continuous stimuli grew with the feature distance between previous and current stimuli. This is in accordance with the adaptation literature, where the previous stimulus with a stronger feature exerts more repulsion on the current stimulus (i.e., adaptation-like repulsion; e.g., Schwiedrzik et al., 2014). In the continuous response task, the strength of previous response and stimulus effects also grew with feature distance between successive events, in accordance with the serial dependence literature (i.e., serial dependence-like attraction). Thus, we have two modulations of attractive previous response effects across response types, one gets stronger and another weaker with increasing relative orientations. However, our design did not include relative orientations between successive stimuli greater than 60°, making it impossible to examine whether the history effects continued increasing or returned to baseline for very large differences; thus, it would be interesting to explore in future studies how history effects behave in a larger extent of feature space. Further studies would be needed to provide a fully mechanistic account, especially given that several other studies found opposing effects with continuous response measures (cf. Moon and Kwon, 2022; Pascucci et al., 2019; Sadil et al., 2024; Zhou et al., 2024). A study with confidence measures comparing response types may be needed for such a mechanistic account.

Our study shows that attractive and repulsive history effects work against each other. In the continuous response task, for example, we found that repulsion from the previous response (in the sequential condition) inhibited the attraction in the condition when stronger repulsion would be expected (i.e., when the previous stimulus was presented longer than the current). This also happened with the repulsion from the previous stimulus in the absence of the previous response (i.e., in the alternating condition). Similarly, with categorical responses, repulsion was stronger in the alternating condition, and attraction was stronger in the sequential condition. These results suggest that whatever the mechanisms of the history effects are, attractive and repulsive forces are ever-present and work against each other, thus arguing for a dynamic visual system that leverages moment-to-moment fluctuations in the visual environment.

Put together, our results reveal the importance of experimental design in determining not only the type of history effects (response vs. stimulus, attractive vs. repulsive) but also the variables modulating these effects. Although our study fails to make a fully mechanistic explanation as to why and how these effects occur, it suggests that independent effects work simultaneously and sometimes against each other in determining behavior. An exciting future path would be to be able to model and discover the neural correlates of history effects between response types.

## Acknowledgements

This work was funded in part by an ANR grant to Thérèse Collins (ANR PRME SEDEP).

## Commercial relationship

None.

